# A mathematical model of Campylobacter dynamics within a broiler flock

**DOI:** 10.1101/574301

**Authors:** Thomas Rawson, Marian Stamp Dawkins, Michael B. Bonsall

## Abstract

Globally, the bacterial genus *Campylobacter* is one of the leading causes of human gastroenteritis, with its primary route of infection being through poultry meat. Despite decades of study we appear to be no closer to preventing outbreaks within commercial chicken flocks, and the application of biosecurity measures is limited by a lack of understanding of the transmission dynamics within a flock. Our work is the first to undertake a mathematical modelling approach to *Campylobacter* population dynamics within a flock of broilers (chickens bred specifically for meat). A system of stochastic differential equations is used to investigate the diverse and fluctuating conditions within the gut of a broiler, and models the routes of infection between co-housed birds. The presented model provides mechanistic explanations for key infection dynamics that have been long-observed but very poorly understood. We highlight several driving mechanisms behind observed infection phenomena, simulate experimentally observed inter-strain competition, and present a promising approach to hypothesising new methods of preventing flock outbreaks.

**Author summary:** The bacteria *Campylobacter* is one of the most common causes of food poisoning globally. The most common route of infection is through raw chicken meat, as a result of many chicken farms across the world housing fully infected flocks. Despite the magnitude of this public health risk, little is understood of the specifics of how chickens become infected, and the ways that they then infect one another. Our work presents a mathematical model of *Campylobacter* transmission dynamics within a flock of chickens. We compare the results of the model to real world data sets, explore key dynamical behaviours, and present a sensitivity analysis to highlight the most important factors underpinning outbreaks.

## Introduction

*Campylobacter* is recognised as the leading cause of human gastroenteritis in the developed world [1]. While several transmission routes have been noted over the years [2], poultry meat has been overwhelmingly attributed as the leading route of ingestion for humans [3]. An ongoing study by Public Health England has highlighted the extent to which *Campylobacter spp.* have dominated our commercial poultry. 73.3% of supermarket chicken carcasses were found to contain *Campylobacter* and 6.8% of the outer packaging was similarly contaminated [4]. An estimated 450,000 people across the United Kingdom are infected every year, with 10% of these infections resulting in hospitalisation [5]. The immediate impact of infection is rarely fatal in the developed world, characterised by stomach cramps and diarrhoea, however the resulting sequelae, while rare, are far more serious. *Campylobacteriosis* leaves the host *∼*100 times more likely to develop the auto-immune disorder Guillain-Barr’e syndrome [6].

While the bacteria provoke an aggressive response in human hosts, the most common species, *Campylobacter jejuni*, is commensal within its most common host, broiler chickens. The term ‘broiler’ refers to any chicken bred and raised specifically for meat production. Once *Campylobacter* is present in a flock, full colonisation of all birds occurs very rapidly [7]. From the introduction of one infected bird, it can take only a single week for an entire flock to become infected [8]. The bacteria are spread via the faecal-oral route. After becoming infected, the newly-infected host broiler spends a brief period in a non-infectious incubation period, before excreting the bacteria in its faecal and cecal matter. Surrounding susceptible broilers are then exposed to this by ingesting the surrounding feed and water [9]. While the direct cause of introduction to the flock is uncertain, an exhaustive review by Adkin et al. (2006) [10] considered that horizontal transmission is by far the most likely route, primarily being brought into a susceptible flock from some other source on the farm, such as the enclosures of other farm animals. This is as opposed to vertical transmission from breeder flocks, which are themselves often fully colonized by *Campylobacter spp.*. Nevertheless, there may be a combination of both routes of entry into a flock, which deserves greater consideration.

*Campylobacter* is very rarely observed to colonise the gut of very young chickens (0 to 2 weeks of age) [11]. This is theorised to be the result of a supply of innate maternal antibodies acquired during a pre-laying period. This immunity has been shown to have significant bactericidal properties [12].

Despite numerous intervention measures being trialled and employed on farms, little impact has been seen in reducing outbreak incidence [13]. This is due in part to the aggressive rate of proliferation once *Campylobacter* has entered a flock, coupled with persisting uncertainty in the exact route of primary infection. Specifically designed prevention methods are also marred by genetic variation and plasticity of *Campylobacter spp.* [14].

Of increasing concern is the growing trend of antimicrobial resistance in *campylobacteriosis* outbreaks. Roughly 90% of the antibiotics applied in agriculture are used only to promote growth or as prophylactic agents, as opposed to being used to treat infection [15]. This overzealous use has been a major contributing factor to the continuing spread of antibiotic resistance. Ge et al. (2003) [16] conducted a study showing that 94% of tested raw chicken samples were resistant to at least one of seven antibiotics being tested, 54% of which showed resistance to erythromycin, the antibiotic most commonly used to treat *campylobacteriosis*. These anti-microbial strains cause more prolonged and severe illness in humans [17] and create a scenario where *in-vitro* susceptibility testing may be necessary before any drugs may be prescribed.

Despite a wealth of empirical investigations, there is a lack of knowledge synthesising these empirical findings through theoretical modelling frameworks. Only two studies have considered a theoretical approach to understanding *Campylobacter spp.* outbreaks; Van Gerwe et al. (2005) [18] and Hartnett et al. (2001) [19], who built a basic SI model and a probabilistic model, respectively. Both frameworks only consider a model on the scale of a flock through basic susceptible-infected interactions. These approaches are not sophisticated enough to develop any meaningful theories on *Campylobacter* dynamics, as they do not represent or convey any specific interbacterial actions by *Campylobacter* populations. The lack of modelling approaches is likely due in part to the inherent challenges of mathematically simulating a gut microbiome. Over 100 different bacterial genera have been isolated from the intestines of chickens [20], all with a range of individual ecological interactions with one-another. Questions must then be asked regarding how to simulate the temporal and spatial impact of gut motility on the development of a microbial community. Despite these challenges, simplified models of stochastic differential equations have proved effective in capturing the often frenetic bacterial population dynamics within the gut [21].

Here, we introduce a framework of stochastic differential equations that captures the basic interactions that are known to be observed within the broiler gut. Using this framework we simulate the propagation of multiple strains of *Campylobacter* through multiple birds in a flock. In the analysis presented below we observe key dynamical behaviour commonly observed through experimentation, which can now be mechanistically explained using this theoretical framework. The theoretical insights derived from this model can be used to refine current hypotheses regarding *Campylobacter* transmission and inform future experimental and control efforts.

## 1 Modelling Frameworks

### 1.1 Deterministic Model

Before presenting the stochastic differential equation framework, we begin by introducing the underlying deterministic core of the framework and the particular interactions modelled. Consider four variables to describe the bacterial populations within a broiler’s digestive tract. *C*, the proportion of a single bird’s gut flora made up of *Campylobacter*. *B*, the proportion of the gut flora made up of other bacterial species competing for space and resources. *P*, the proportion of the gut containing host defence peptides (HDPs) (this may also be interpreted as other plausible forms of host autoimmune response). Lastly, *M*, the proportion of the gut containing innate maternal antibodies. These all take values ranging such that 0 ≤ *C*, *B*, *P*, *M* ≤ 1. The set of ODEs describing the dynamics follows:

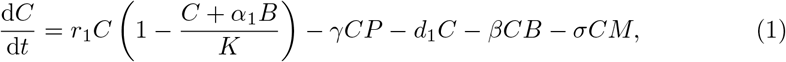

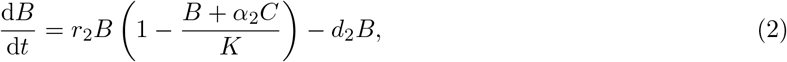

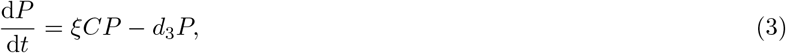

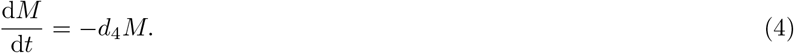

All rate constants are defined below in Table 1. The first term 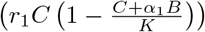 in equation (1) describes the logistic growth of *Campylobacter* to a carrying capacity, *K*, while in competition with other bacteria *B*. Competition for resources is the key to success within the gut. *Campylobacter* is known to be an effective coloniser [22], as it is very effective at drawing zinc [23] and iron [24] from its environment. The second term (*γCP*) in equation (1) models the inhibitory effect of host defence peptides, *P*. These peptides are created in response to challenge by *Campylobacter*, as shown by Cawthraw et al. (1994) [25]. The third term (*d*_1_*C*) of equation (1) simply describes the natural death rate of *Campylobacter*. The fourth term (*βCB*) simulates an important interbacterial interaction; that some of the most abundant competing bacteria in the microbiome have an inhibitory effect on *Campylobacter* [26]. The final term (*σCM*) of equation (1) represents the strong bactericidal abilities of the bird’s maternal antibodies. All chickens hatch with an initial supply of antibodies that depletes over time, gone by about three weeks of age [12] (most broilers are slaughtered at five or six weeks of age, however some organic and free-range flocks are slaughtered at approximately eight weeks). These antibodies have a strong inhibitory effect on *Campylobacter*, and many studies are unable to detect *Campylobacter* (by culture methods) in birds under 2 weeks of age under commercial conditions [27]. However, forced inoculation of high-quantities of *Campylobacter* soon after hatching can still result in expression of the bacteria [28].

**Table 1.** Model parameters and baseline values. Descriptions for all parameter values appearing in the final stochastic model, equations (14) - (18). Baseline values are given, used for model validation and simulation case studies. *Ω value is dependent on the experiment specifics for model validation, but flock case studies consider a flock of 400 chickens, and an Ω value of 200,000.

| Expression | Description | Value |
| --- | --- | --- |
| $r_{C_j}$ | Growth rate for *Campylobacter* strain $j$. | 0.27 |
| $r_2$ | Growth rate for other bacteria ($B$). | 0.15 |
| $\alpha_1$ | *Campylobacter* competition coefficient. | 0.92 |
| $\alpha_2$ | Other bacteria competition coefficient. | 1 |
| $K$ | Carrying capacity. | 1 |
| $\gamma_{C_j}$ | Rate of inhibition by host defence peptides ($P$) on *Campylobacter* strain $j$. | 0.2 |
| $\xi_j$ | Rate of host defence peptide growth in response to *Campylobacter* strain $j$. | 0.4 |
| $b$ | Rate of broiler shedding *Campylobacter* into the environment, $E_j$. | 10 |
| $\Omega$ | Total environmental carrying capacity of *Campylobacter*. | 200,000* |
| $d_{C_j}$ | Death rate of *Campylobacter* strain $j$. | 0.02 |
| $d_2$ | Death rate of other bacteria. | 0.02 |
| $d_3$ | Decay rate of host defence peptides. | 0.05 |
| $d_4$ | Decay rate of maternal antibodies. | 0.005 |
| $d_5$ | Death rate of *Campylobacter* in the environment. | 0.05 |
| $\beta_{C_j}$ | Rate of inhibition by other bacteria on *Campylobacter* strain $j$. | 0.03 |
| $\sigma_{C_j}$ | Rate of inhibition by maternal antibodies on *Campylobacter* strain $j$. | 0.07 |
| $\eta_{C_j}$ | Scaling factor applied to stochastic *Campylobacter* growth in the gut. | 0.01 |
| $\eta_{BC_j}$ | Scaling factor applied to stochastic *Campylobacter* inhibition by other competing bacteria. | 0.09 |
| $\eta_2$ | Scaling factor applied to stochastic competing bacteria ($B$) growth. | 0.01 |
| $\eta_3$ | Scaling factor applied to stochastic host defence peptide ($P$) growth. | 0.01 |
| $\eta_4$ | Scaling factor applied to stochastic maternal antibody ($M$) decay. | 0.01 |
| $\eta_5$ | Scaling factor applied to stochastic *Campylobacter* growth in the environment. | 0.01 |

Equations (2), (3) and (4) follow a similar logic to equation (1). Other bacteria, *B*, grow in competition with *Campylobacter* to a carrying capacity. Defence peptides, *P*, grow in response to *Campylobacter* expression (not in competition for resources), and the population of maternal antibodies, *M*, does not grow. All variables decay at a rate proportional to their respective populations.

Note that the above model could be reduced by amalgamating terms in equations (1) and (2), however we choose to keep these separate to (i) keep biological processes clearly defined, and (ii) make further model development and sensitivity analyses clearer.

Ignoring the trivial cases of complete domination by either *C* or *B*, the basic dynamical behaviour observed for this simplified model is illustrated in Figure 1. Notably, *Campylobacter* is absent from the microbiome until the maternal antibody population has been exhausted. At this point a sudden, temporary, surge in the population of *Campylobacter* is observed. This phenomena is due to the very low population of HDPs, caused by the strong effect of the initial maternal antibodies. The HDP population then quickly rises to meet this sudden challenge, bringing the *Campylobacter* population back to a lower level in an oscillating manner, where it eventually reaches a steady-state equilibrium. This behaviour is commonly observed in experimental studies [29] [30].

**Fig 1.**
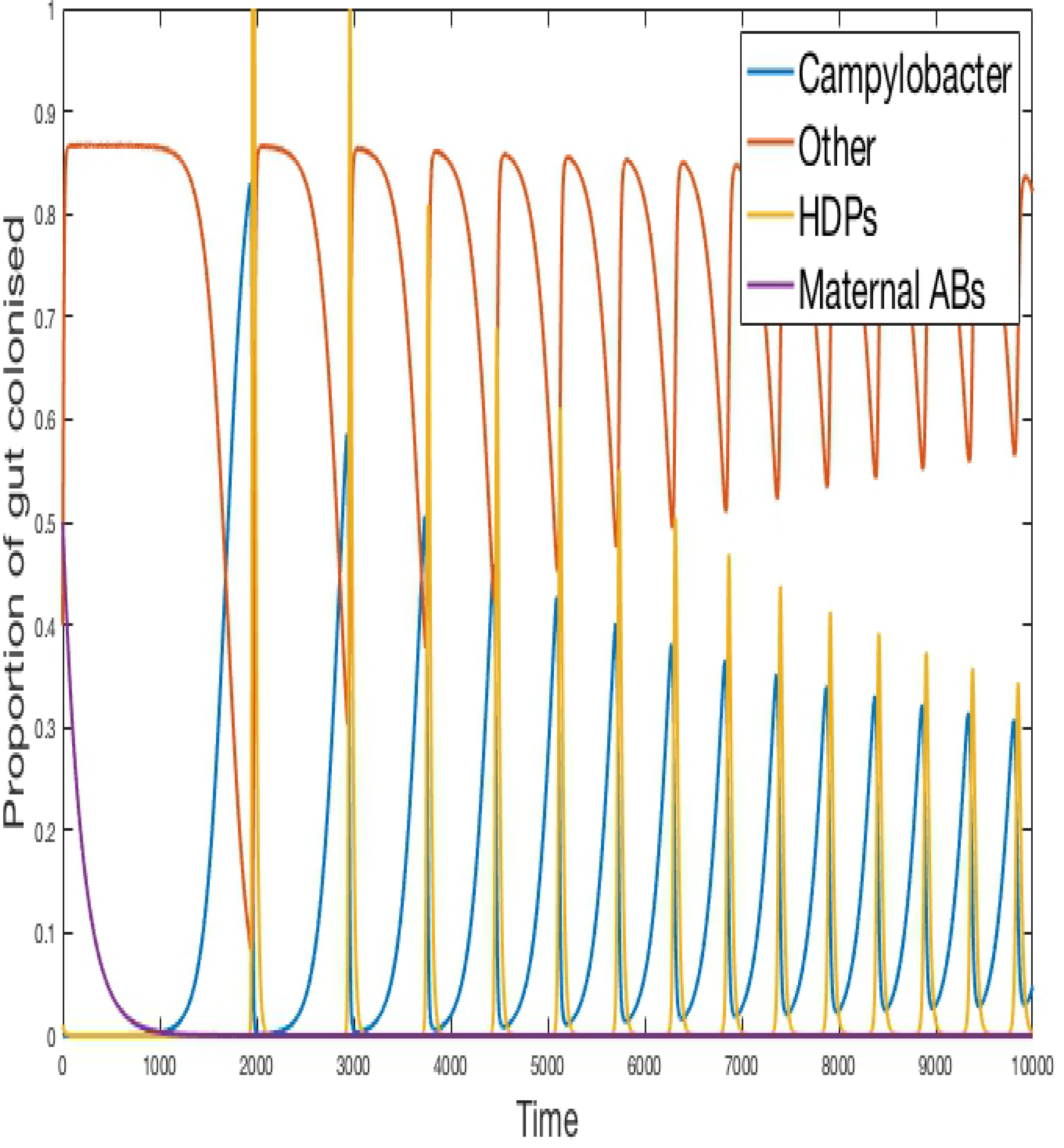
Deterministic model for one chicken. An example of the typical dynamical behaviour observed for simulations of equations (1) - (4). Parameters defined in Table 1.

From this simple core of four equations we adapt the model to allow for *N* unique strains of *Campylobacter*, by describing each strain as a separate variable. Equation (1) is repeated for each individual strain, while altering the growth rate terms to reflect the fact that all strains will also be in competition with one another. This alteration is represented by the following set of ODEs:

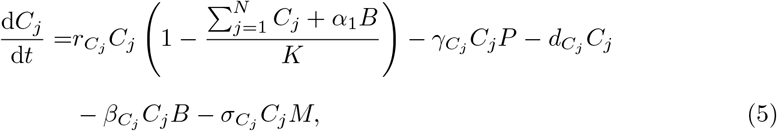

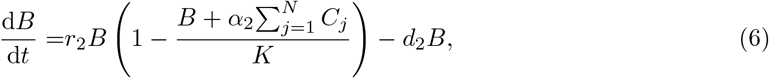

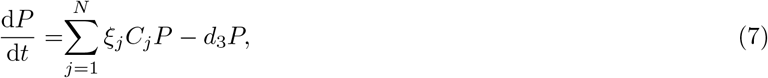

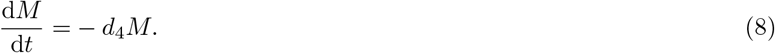

Here *C*_*j*_ represents the *j*^th^ strain of *Campylobacter*, where *j* ∈ {1, 2, …, *N*}, and *N* is the total number of strains. As such this adjusted model is composed of *N* + 3 variables. The next alteration is to allow for multiple birds and the ability for *Campylobacter* to move from one bird to another. This is done by repeating the *N* + 3 equations presented in equations (5)-(8) for each bird, and introducing new variables to display the saturation of *Campylobacter* strains in the shared living space.

As such, the newly adjusted model to describe the population dynamics of *N* strains of *Campylobacter* within *L* broilers, is written as,

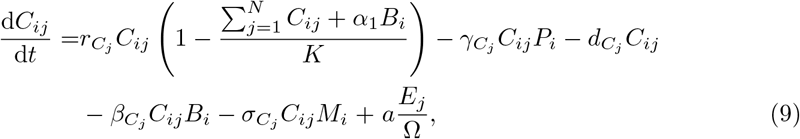

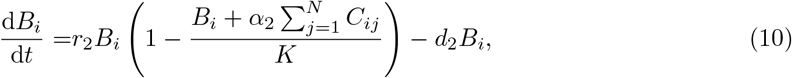

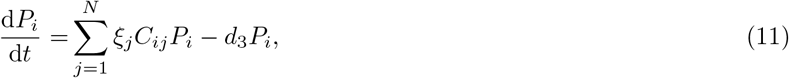

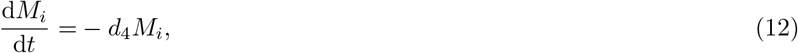

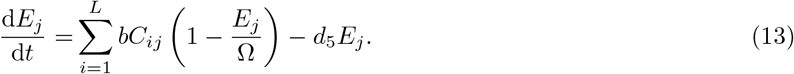

Here then, *C*_*ij*_ represents the proportion of the *i*^th^ broiler’s gut bacteria which is composed of *Campylobacter* strain *j*. *B*_*i*_ is the proportion of the *i*^th^ broiler’s gut bacteria made up of other bacterial species competing for space and resources. *P*_*i*_, the proportion of the *i*^th^ broiler’s gut containing host defence peptides. *M*_*i*_ is the proportion of the *i*^th^ broiler’s gut containing innate maternal antibodies. Here *i* ∈ {1, 2, …, *L*}, where *L* is the total number of broilers. *E*_*j*_ represents the amount of *Campylobacter* strain *j* that is currently in the flock’s enclosed living space. We assume a living space of fixed size shared by all broilers. As such, Ω represents this total size, or carrying capacity for strains. The first term in equation (13) shows that the amount of strain *j* in the environment is increased by being shed from birds that are already infected with strain *j* at a rate *b*. Note from the final term 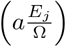 in equation (9) that birds may then ingest strain *j* from the environment at a rate 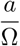. This route of infection simulates the faecal-oral route of infection, but may be interpreted as some other intermittent transmission stage between birds. The model is now composed of *L*(*N* + 3) + *N* equations, for *N* strains of *Campylobacter*, and *L* individual broilers.

### 1.2 Stochastic Model

While several important biological phenomena can be discovered and better understood with the model in its current, deterministic, form, there are key reasons to pursue a stochastic framework. First, having one variable alone to represent the multitudes of bacterial species that make up the constantly-evolving gut microbiome is, of course, a significant simplification. In practise, these other bacterial species competing with *Campylobacter* will be constantly changing, both in resurgences of population and in how they interact with *Campylobacter*. Adding stochastic elements to these populations and interactions is a small step towards capturing some of this more unpredictable behaviour. Indeed the biomass of *Campylobacter* measurable in faecal and cecal matter has been observed to fluctuate widely [29] [31]. Secondly, the law of mass action assumptions made when formulating the initial deterministic model are assumptions that break down for smaller populations. The simulations undertaken often display bacterial populations at very small quantities, especially in the initial period dominated by maternal antibodies. A stochastic system behaves very differently under these circumstances and means that the model is more likely to display cases of strain extinction, a phenomena that the deterministic model cannot capture. Indeed, the very nature of *Campylobacter* infections is one that is often described in the language of probability. The all-or-nothing nature of flock infections means that we often must ask what measures can reduce the likelihoods of infections, rather than the magnitude. Through a stochastic framework we explore multiple realisations of potential outcomes, and investigate reducing the likelihood of outbreaks.

For the stochastic framework, equations (9)-(13) are adjusted to the following set of stochastic differential equations,

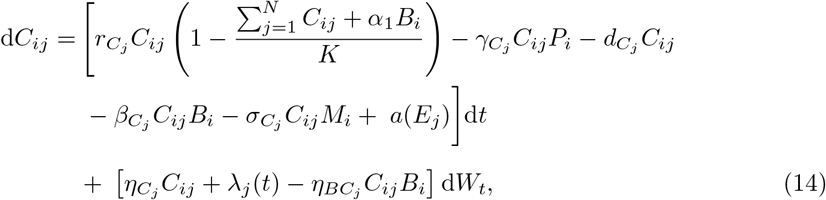

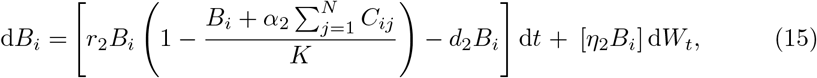

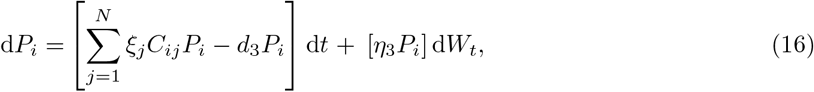

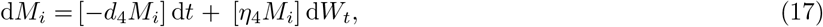

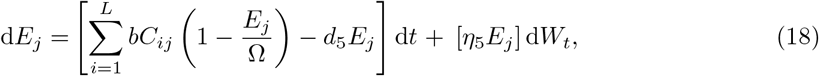

where *λ*_*j*_(*t*) is defined by;

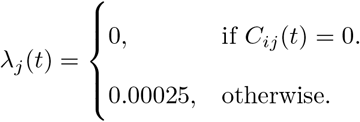

and where *a*(*E*_*j*_) is defined by;

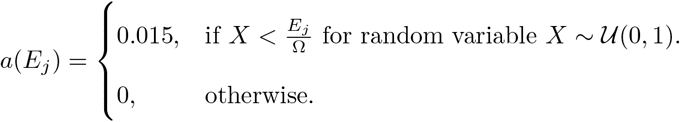

The stochastic additions in equations (15) - (18) are a Wiener process applied to the population (standard Brownian motion process), scaled by the respective population size and constants *η*_2_ through to *η*_5_. These constants dictate the variance of their respective Wiener processes, defining the range of stochasticity attributed to the growth rate of their respective variables. The changes and additions shown in equation (14) warrant further explanation. The sixth term (*a*(*E*_*j*_)) in equation (14) (the last of the deterministic terms), has been changed from a constant rate of ingestion from the environment, as seen in equation (9), to instead have ingestion modelled by a chance to ingest *Campylobacter* depending on the amount of that strain in the environment, *E*_*j*_. The greater *E*_*j*_ is, the more likely it is for ingestion to occur.

The eighth term (*λ*_*j*_(*t*)) in equation (14) is a Wiener process independent of the population of *C*_*ij*_. This is introduced to allow for the possibility of extinction events, should the population of *C*_*ij*_ reach a particularly low threshold. This threshold is decided by the value taken by *λ*_*j*_(*t*), in this case 0.00025. Finally, the ninth term of equation (14) applies a Wiener process around the interactions between *C*_*ij*_ and the competing bacteria *B*_*i*_. This term allows for instances when the particular biodiversity and spatial structure of the gut microbiome may be more inhibitory towards *Campylobacter*, or perhaps actually assisting its growth instead.

Several interesting dynamical behaviours can be observed using this model, which are highlighted through some specific question-led case studies. Table 1 defines all parameters presented in the final stochastic model ((14) - (18)) as well as a baseline of parameter values that were used in model validation against real world data sets (presented below). The model is constructed to an arbitrary timescale, however the parameter values given in Table 1 ensure that multiple oscillations in the *Campylobacter* population can be observed in the below case studies, a phenomena observed in the lifespan of broilers [31]. Broilers are usually slaughtered at approximately five weeks of age, and maternal antibodies (*M*) are usually depleted after approximately three weeks.

Note that throughout we have chosen to use a *Campylobacter* competition coefficient of *α*_1_ = 0.92 < 1. This choice is justified in that bacterial populations can inhabit multiple intestinal niches that cannot be colonised by other competing bacteria. Indeed competitive exclusion therapies have been far less effective in tackling *Campylobacter* compared to other foodborne illnesses such as *Salmonella* [32]. The deterministic model is solved using the ode45 solver, a fifth-order Runge-Kutta method in Matlab. The stochastic model is solved numerically using the Euler-Maruyama method [33] with *N* = 2^14^ timesteps, also programmed in Matlab. The code used to produce all figures presented is available at: https://osf.io/b3duc/.

### 1.3 Model Validation

We test our model by comparing its predictions against three experimental studies on *Campylobacter* expression and spread. Firstly, we consider the work of Achen et al. (1998) [29]. Achen et al. performed an experiment with twenty-four broilers, who were kept in individual, isolated wire-bottomed cages. Birds were confirmed as free of *Campylobacter* before being inoculated with a *C. jejuni* suspension. A cloacal swab was then obtained from each bird every day for forty two days, to monitor whether or not each bird was shedding *Campylobacter*. Figure 2 shows their experimental results alongside the predictions made by our model.

**Fig 2.**
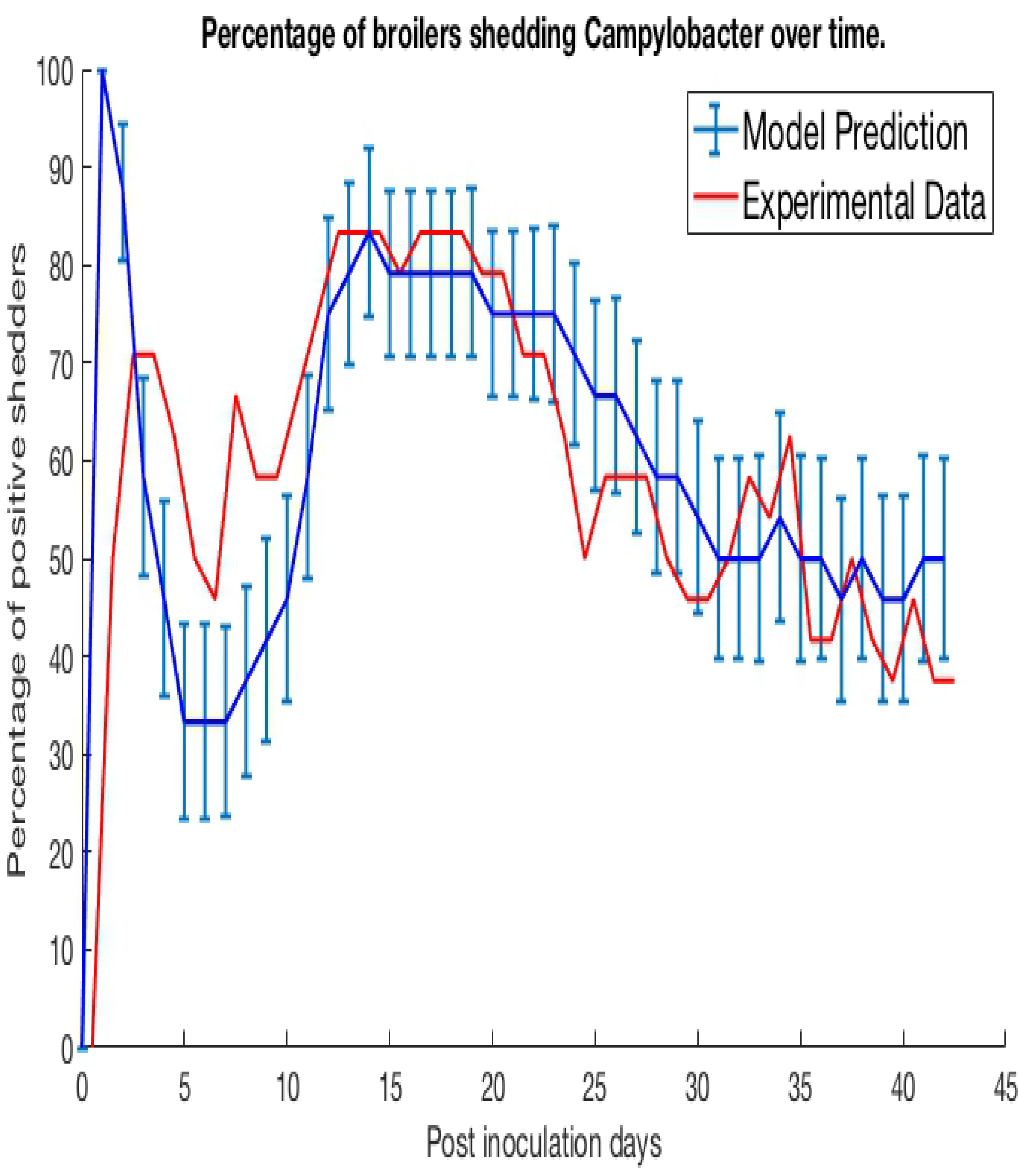
Model validation against data of Achen et al. (1998) [29]. A graph plotting the percentage of a group of isolated broilers shedding *Campylobacter* across several weeks following inoculation.

Specifically, the blue line represents the modal value of the percentage of the 24 birds shedding across a thousand simulations, with error bars depicting the standard deviation across these simulations. Achen et al. (1998) also reports how most birds would shift from phases of positive shedding to negative shedding, a phenomena also captured by the oscillating behaviour displayed by the model. Sampling via culture methods like those performed in this experiment is prone to false-negative results for samples with very low quantities of *Campylobacter* [34]. Therefore, for this model validation, we considered a broiler as being clear of *Campylobacter* if its proportion of *Campylobacter* (variable *C*) was below 0.005. This was considered a more accurate measure to correspond with the experimental data. While our model was constructed to an arbitrary timescale, comparing to this real-world data set it was found that our timescale is approximately equal to *t* = 1 *∼* 30 minutes.

Secondly, we consider the experiment conducted by Stern et al. (2001) [8]. Multiple separates pens were prepared, each containing 70 broilers, all free of *Campylobacter*. A *Campylobacter*-positive seeder bird was then added to the flock. Different pens had seeder birds introduced at different points in time. 3, 5 and 7 days after a seeder bird was introduced, a sample of chickens were tested for *Campylobacter* to estimate the percentage of the flock that was currently *Campylobacter*-positive. We plot our model predictions against Stern et al.’s (2001) experimental data below in Figure 3. To match the housing density of the experiment, a value of Ω = 45, 369 was used for the model. An error band is plotted around our model prediction displaying the standard deviation of values across 100 simulations.

**Fig 3.**
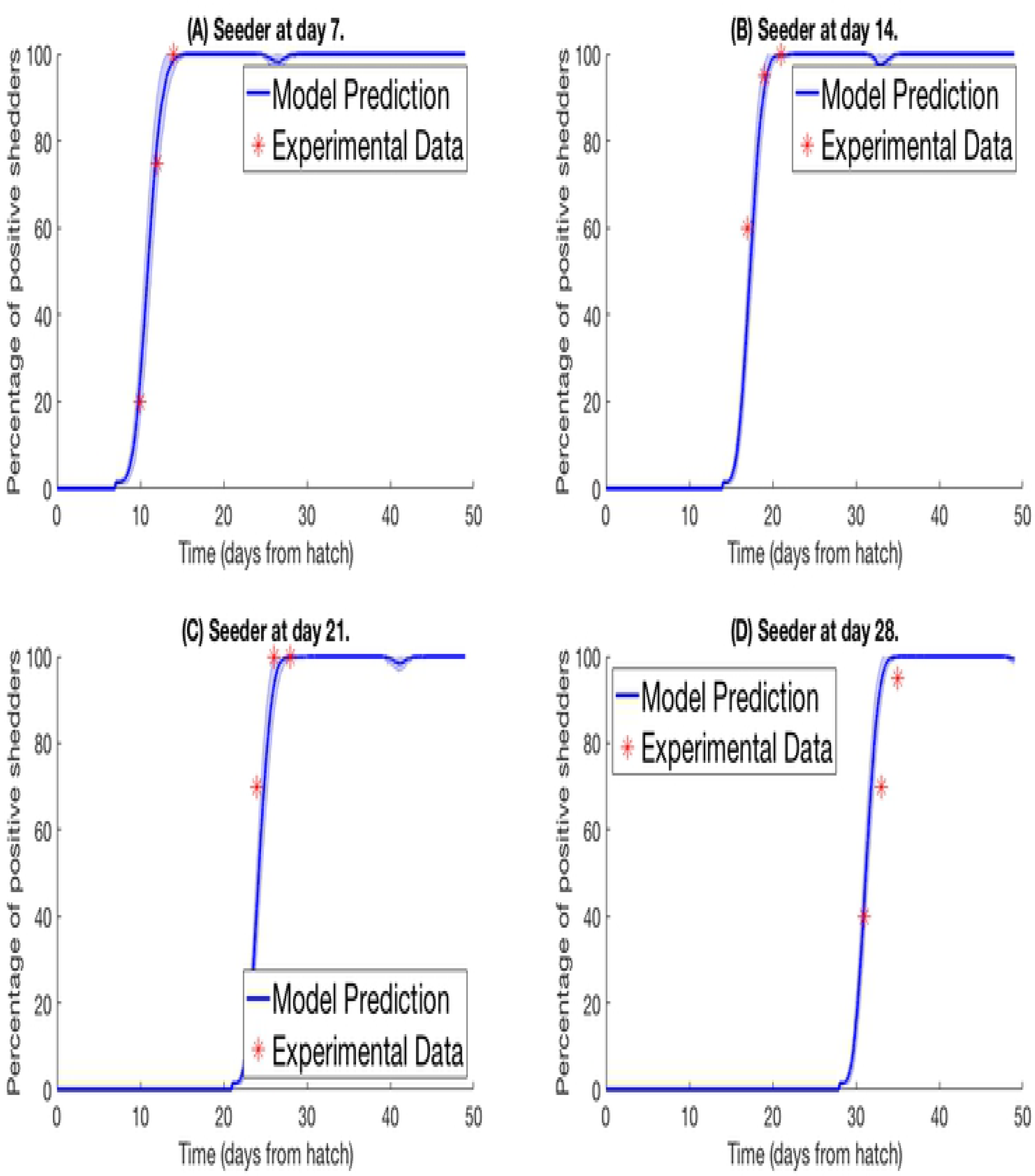
Model validation against data of Stern et al. (2001) [8]. Graphs plotting the percentage of a lock of broilers shedding *Campylobacter* across several weeks after introduction of a *Campylobacter*-positive seeder bird at **(A)** seven days, **(B)** fourteen days, **(C)** twenty one days, **(D)** twenty eight days.

Lastly we simulated the experiment performed by Van Gerwe et al. (2005) [18]. Four flocks of 400 birds were set up in individual enclosures from day of hatch. Four birds in each flock were then inoculated with a *Campylobacter* suspension and returned to the flock. Birds were then sampled from each flock throughout the next few weeks to record the percentage of flock infection. Figure 4 plots their experimental data against our model prediction. For the experiments shown in Figure 4A and Figure 4B, the four seeder birds were inoculated at day of hatch, and chickens were sampled by cloacal swabbing. For the experiments shown in Figure 4C and Figure 4D, the seeder birds were inoculated one day after hatch, and the flock was analysed by collecting fresh fecal samples.

**Fig 4.**
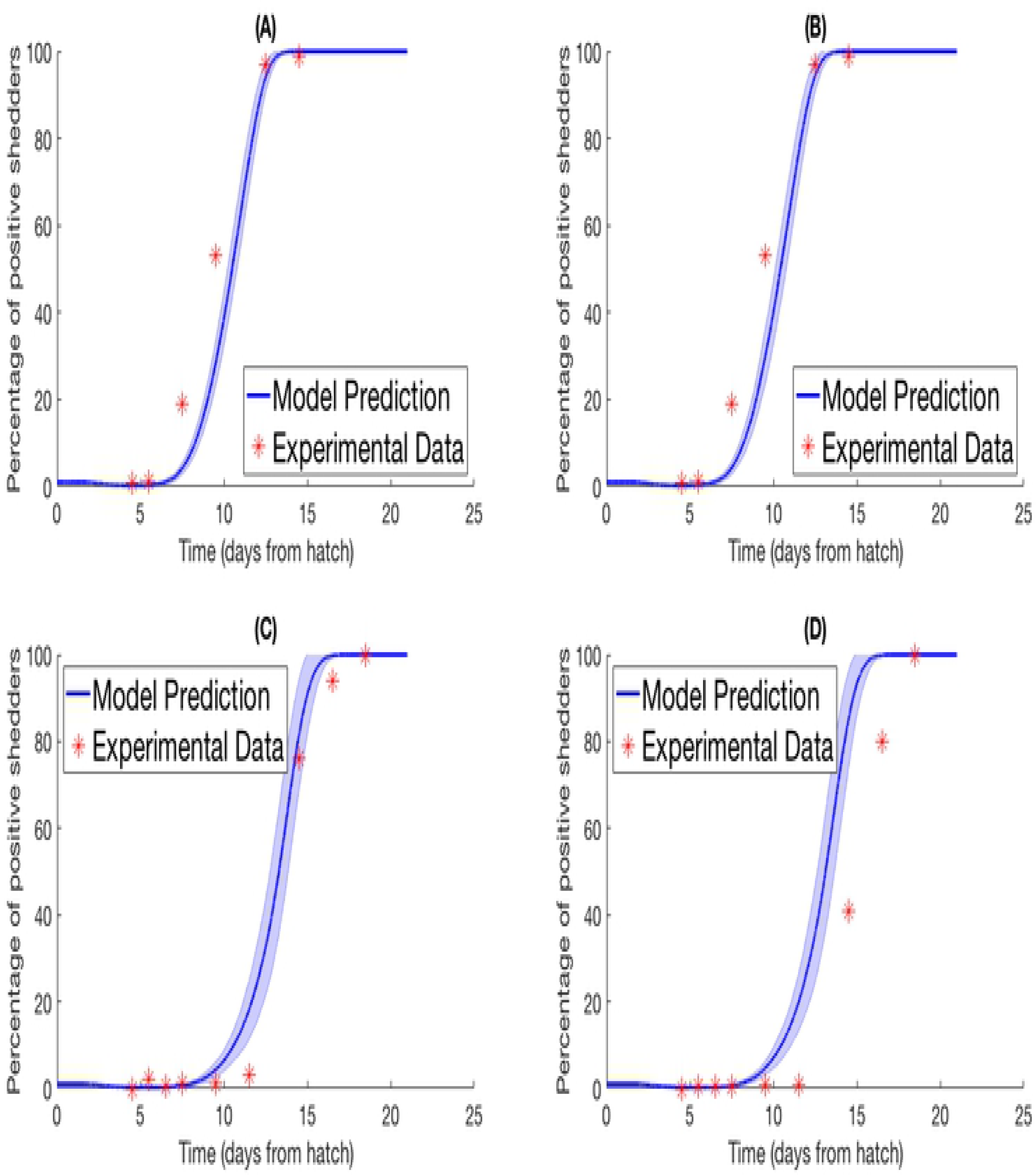
Model validation against data of Van Gerwe et al. (2005) [18]. Graphs plotting the percentage of a flock of broilers shedding *Campylobacter* across several weeks after introduction of a *Campylobacter*-positive seeder bird. (A)/(B) Seeder bird introduced at day of hatch, samples collected via cloacal swab, (C)/(D) seeder bird introduced one day after hatch, samples collected via fresh fecal droppings.

## 2 Simulations

We now use a series of (simulated) case studies to investigate key dynamical behaviours and predictions from the model.

### 2.1 Staggered Strain Infection

In this first example, the deterministic model for multiple strains in one broiler (equations (5) - (8)) is considered. Five strains of *Campylobacter* within one chicken are simulated, all with the exact same respective rate constants as shown in Table 1. Figure 5A shows the results when all five strains are introduced at *t* = 0 with the same initial inoculation amount of *C*_*i*_(0) = 0.0001. Figure 5B shows instead when each strain is introduced in intervals of *t* = 250. Therefore only strain 1 is introduced at *t* = 0, strain 2 is introduced at *t* = 250 and so on until finally strain 5 is introduced at *t* = 1000. In both cases the other three variables are initialised at *B*(0) = 0.4, *P* (0) = 0.01 and *M* (0) = 0.5.

**Fig 5.**
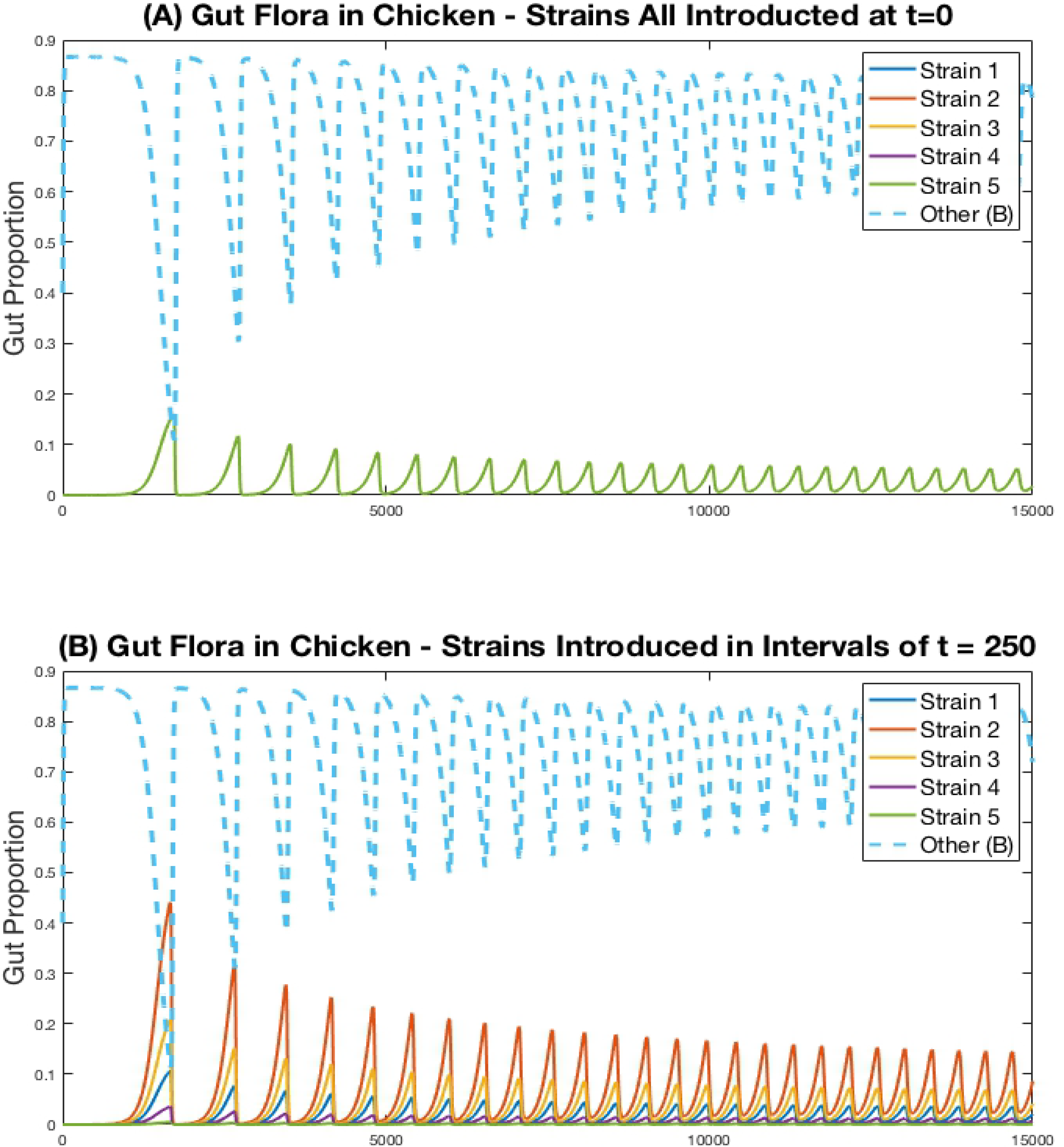
Simulations of multiple *Campylobacter* strains within one broiler. Population growth of five strains of *Campylobacter* within one broiler that are **(A)** all introduced at *t* = 0 **(B)** introduced in intervals of *t* = 250. Strains are initialised at *C*_*j*_(*t*) = 0.0001 at their respective time of introduction. Other variables are initialised at *B*(0) = 0.4, *P* (0) = 0.01 and *M* (0) = 0.5. Note that the single green line in Figure 5A is due to overlap, all five strains exhibit the exact same dynamical behaviour, as would be expected.

While the maternal antibodies (*M*) are not plotted on these figures, they approach 0 at approximately *t* = 1, 000, as can be seen by the following surge in *Campylobacter* populations following this point in Figure 5. While, unsurprisingly, all strains perform identically in figure 5A (where strains are initialised at the same point in time), a more curious dynamic is observed in Figure 5B. The strain that performs best and exists at the highest proportion in the staggered release example is strain 2, the second strain to be introduced. The reason for this is that strain 1, present at *t* = 0, is initially suppressed by the maternal antibodies (parameter *M*), reducing the proportion of strain 1. As a result, when strain 2 is introduced, it is able to capitalise on the severely reduced amount of strain 1, and the reduced amount of maternal antibodies, to quickly grow and dominate the competitive space. Strain 2’s increased presence then puts future strains at a disadvantage as it has already had the opportunity to establish itself within the gut. These results suggest that dominant *Campylobacter* strains can prevent new strains from taking hold. Moreover, there is an optimal point in time for inoculation to occur for a strain to become dominant, as shown in Figure 5B where strain 2 is consistently occupying a higher proportion of the gut than other strains.

### 2.2 Stochastic model - One strain in one broiler

The stochastic model (equations (14) - (17)) is run to simulate one strain of *Campylobacter* within one broiler. In this scenario, we ignore the environmental variable *E* (equation (18)), as its input is negligible for only one broiler. The rate constants are kept at the same values as used previously, defined in Table 1, with the additions of the stochastic variance scaling rate constants, parameters that limit the variance of the stochastic additions. These are set as *η*_*Cj*_ = *η*_2_ = *η*_3_ = *η*_4_ = 0.01, and *η*_*BCj*_ = 0.09. *η*_*BCj*_ is set higher than the other stochastic rate constants to capture the greater unpredictability surrounding these bacterial interactions. Four different realisations of this model are presented in Figure 6, all initialised at *C*(0) = 0.02, *B*(0) = 0.4, *P* (0) = 0.01, *M* (0) = 0.5.

**Fig 6.**
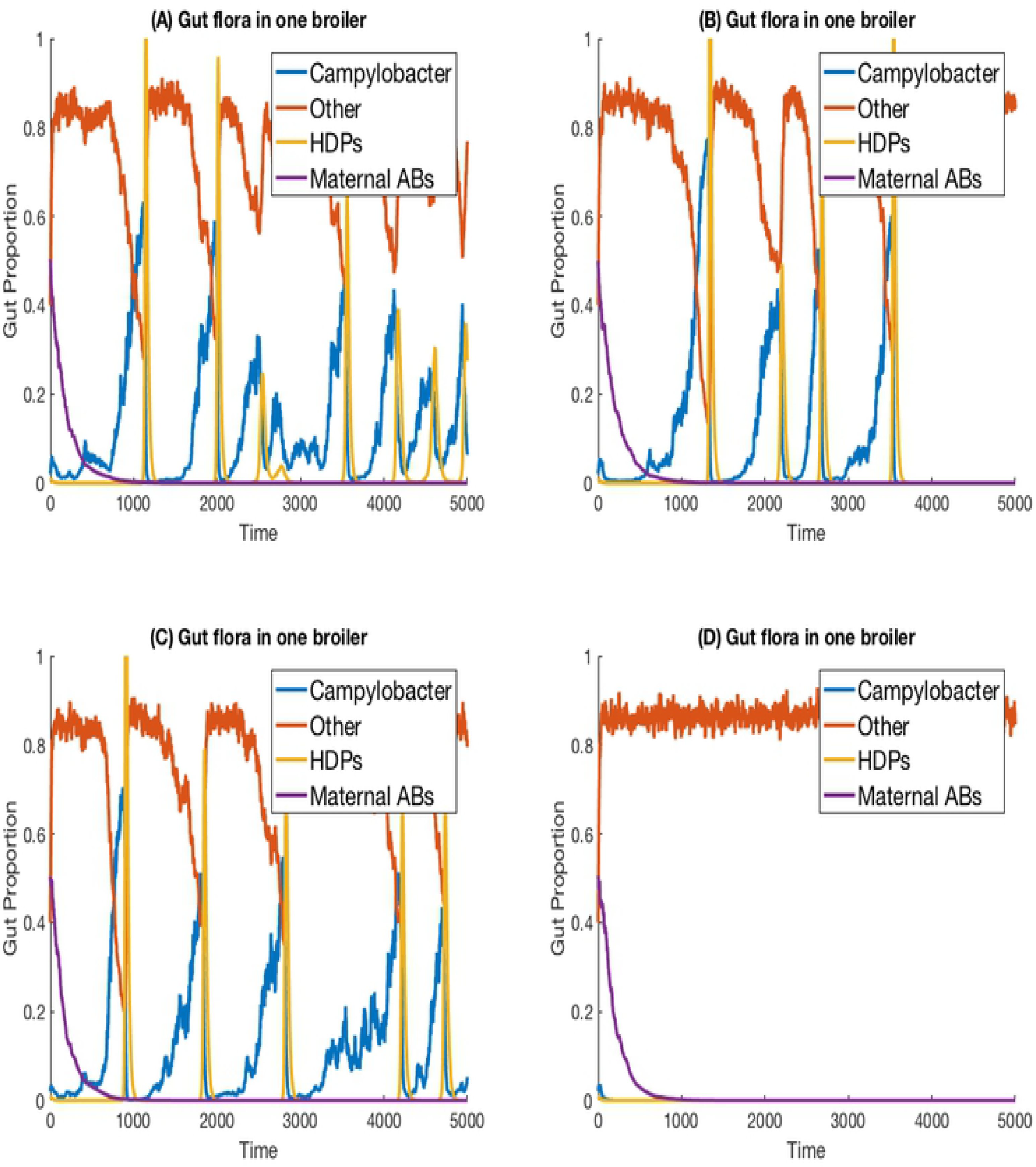
Stochastic simulations of one *Campylobacter* strain within one broiler. Four different realisations of a stochastic model simulating one strain of *Campylobacter* within one isolated broiler.

Empirical studies measuring the amount of *Campylobacter* in the faecal matter of isolated broilers have shown a spectrum of results. Some broilers display sustained high populations, others express initial peaks followed by great reduction and potentially later resurgence, and sometimes extinction cases are observed [29]. All these dynamical behaviours can be observed in different realisations of this model (Figure 6). Figure 6A shows an instance where a broiler is consistently infected and shedding into the environment, unable to effectively clear the *Campylobacter* from its system. Figure 6B instead shows an instance where a broiler has multiple periods of high infection and shedding, before being able to clear the infection. Figure 6C shows similar behaviour to 6A, whereby the broiler is unable to clear the bacteria, however 6C shows more dramatic peaks and troughs in its dynamic profile, suggesting it may have longer periods of reduced shedding. Finally, Figure 6D shows an instance where the broiler successfully clears *Campylobacter* at the initial point of inoculation. All these realisations are run with the same parameters given in Table 1, demonstrating the benefit of a stochastic framework being able to better capture the more diverse range of possible events.

### 2.3 Stochastic model - One strain in multiple broilers

The previous scenario is now extended to consider multiple broilers. Figure 7 presents the results for one *Campylobacter* strain in a flock of 400 broilers. We use the parameter values stated in Table 1. The total size of the enclosure, or the carrying capacity of *E*, is set at Ω = 200, 000. This value is considered in cm^2^, and so with 400 broilers, translates to 500cm^2^ per broiler. EU directive 2007/43/CE states that broilers may never be stocked at more than 42kg/m^2^ [35]. Assuming a targeted bird weight of 1.5kg, this translates to 357cm^2^ per bird. This simulation models slightly more space allowed to each bird than the limit. The death rate of *Campylobacter* in the environment is set at *d*_5_ = 0.05, higher than the death rate within a broiler as, despite their many survival mechanisms [36] *Campylobacter* is susceptible to many exterior environmental stresses [37] and is exceptionally fragile outside of its host. The simulation began with no *Campylobacter* in the surrounding environment (*E*(0) = 0) and the other initial conditions are set the same as for the previous example, with the exception that two of the 400 broilers start with an initial condition of *C*_1_(0) = *C*_2_(0) = 0.02, while the others are initialised without any *Campylobacter*. These results are shown in Figure 7.

**Fig 7.**
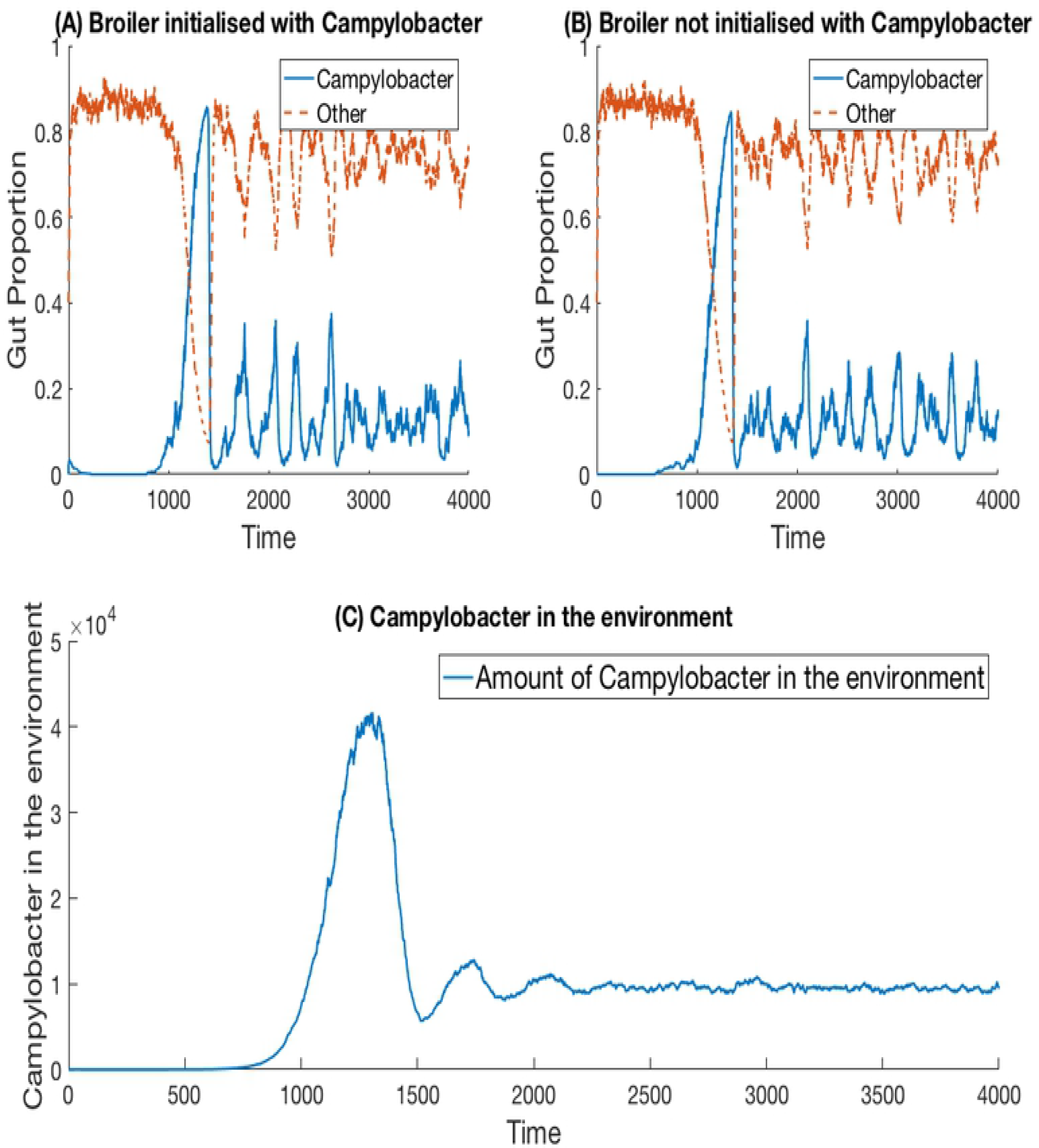
Stochastic simulations of one *Campylobacter* strain within multiple broilers. The proportion of a broiler’s gut containing *Campylobacter* for **(A)** a broiler in the flock initialised with a small proportion of *Campylobacter* **(B)** a broiler in the flock initialised with no *Campylobacter*. **(C)** shows how much of the environment (total size of 200,000) contains *Campylobacter*. This is variable *E* in the model.

While birds who are not initialised with *Campylobacter* become infected at a slightly later time, the dynamical behaviour is very similar across all birds in the flock. Multiple realisations do not display the broader spectrum of behaviour observed in the one broiler case (Figure 6). The implication is that housing a greater number of birds causes more homogeneous dynamical behaviour, and indeed the wide variety of *Campylobacter* expression seen in the isolated bird experiments of Achen et al. (1998) [29] is not so commonly observed in experiments with group-housed birds [18].

### 2.4 Stochastic model - Five strains in multiple broilers

We extend the previous scenario to investigate dynamics of multiple strains. Five strains of competing *Campylobacter* are modelled within the same flock of 400 birds. The same constants are used as in the previous scenario, with each strain having identical rate constants. One key difference is that all broilers are initialised without any *Campylobacter*, instead an initial amount is present in the environment. Each strain of *Campylobacter* in the environment is initialised at *E*_1_(0) = *E*_2_(0) = *E*_3_(0) = *E*_4_(0) = *E*_5_(0) = 100. The results of this simulation are shown in Figure 8.

**Fig 8.**
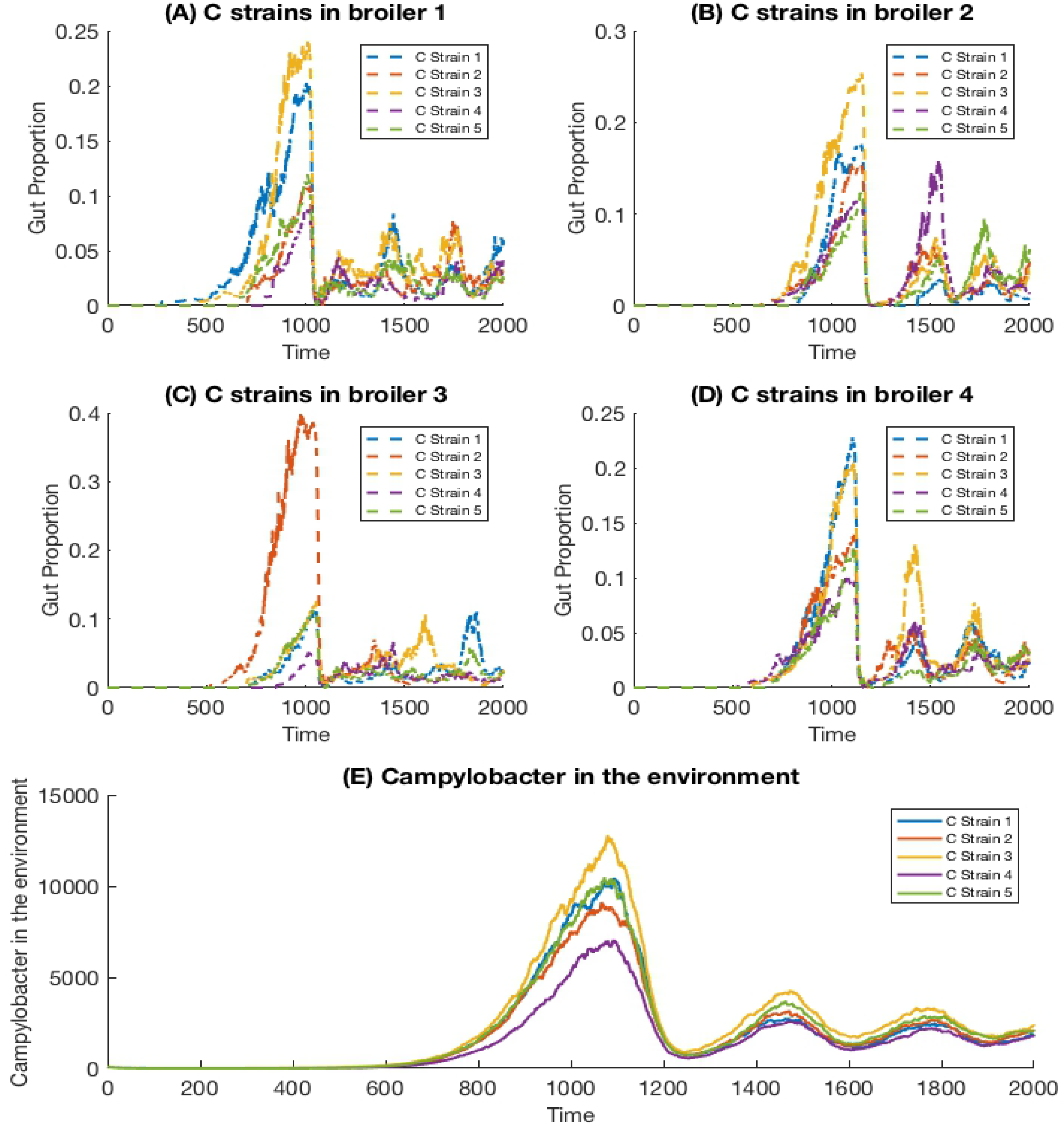
Stochastic simulations of multiple *Campylobacter* strains within multiple broilers. The proportion of four different broilers’ microbiomes that contain five strains of *Campylobacter*. All birds are within the same flock. **(E)** shows how much of the environment (total size of 200,000) contains the five strains of *Campylobacter*. These are variables *E*_*j*_ in the model.

On average, all strains perform equally well across the flock, as shown in Figure 8E. All strains are present at roughly equal amounts in the environment, reflecting an equal presence on average across all birds in the flock. However, when observing the *Campylobacter* proportions within individual broilers, one or two strains will tend to dominate early on in colonising a broiler’s gut, which can in turn prevent other strains from taking hold (seen most clearly in Figure 8C). This dynamical behaviour was first observed in our deterministic simulations (see Figure 5B), however unlike in the deterministic case, stochastic events can cause dominant *Campylobacter* strains to reduce in population, presenting an opportunity for a different strain to establish itself.

This phenomena is more clearly seen if the timescale of the simulation is extended, as illustrated in Figure 9. Although the average population of strains across the flock is equal, the stochastic model shows that a single strain of *Campylobacter* tends to dominate the gut of individual broilers at any one time. Although there are brief periods where strains exist in equal amounts, eventually the balance shifts again to longer periods of dominance by one or perhaps two strains.

**Fig 9.**
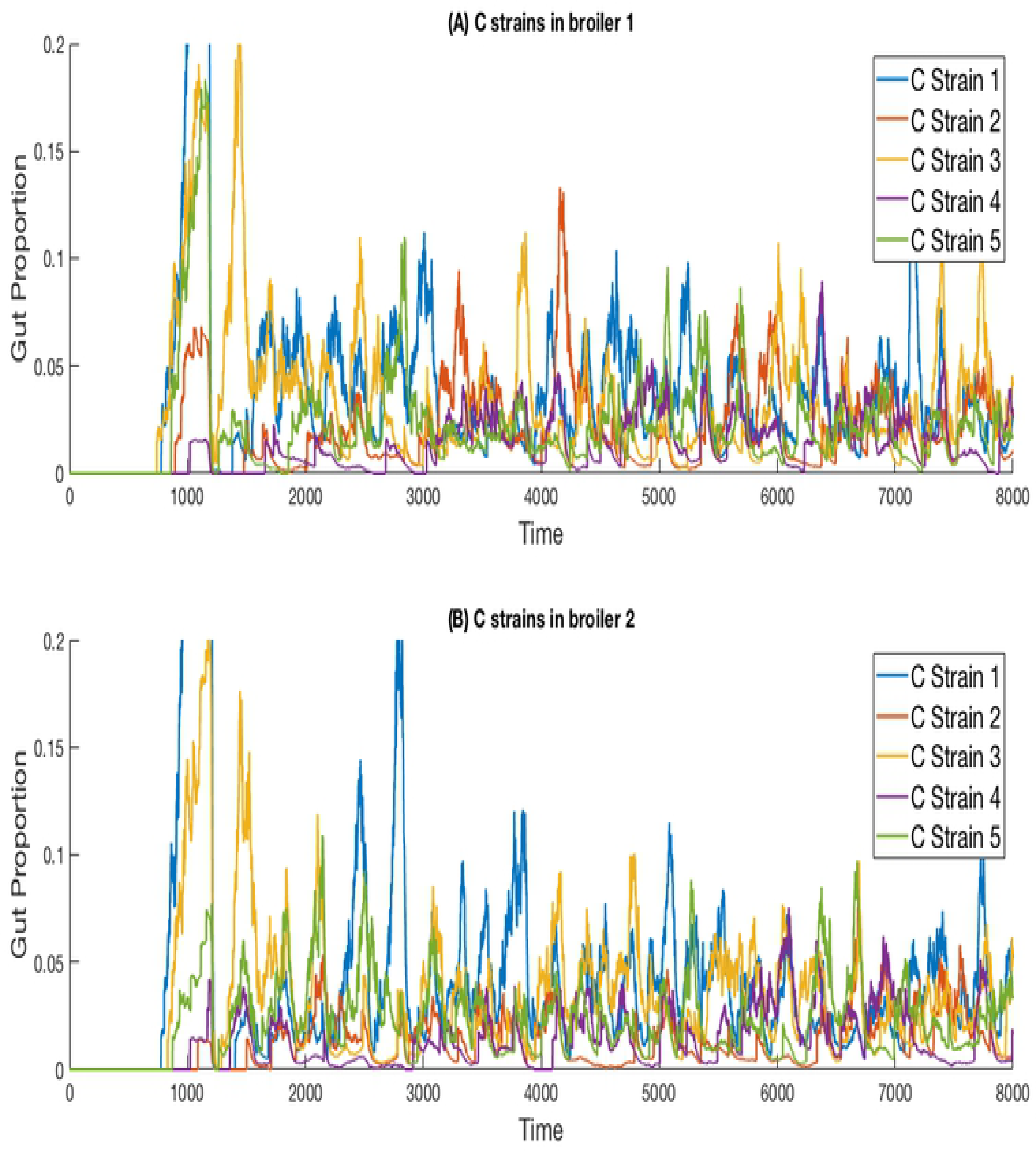
Stochastic simulations of multiple *Campylobacter* strains within multiple broilers across a greater timescale. The proportion of two different broilers’ microbiomes that contain five identical strains of *Campylobacter*.

Disadvantaged strains of *Campylobacter* are quickly eliminated. Figure 10 shows the results for a simulation where strain 4’s growth rate, *r*_*C*4_, is reduced from 0.27 to 0.265, and strain 5’s growth rate, *r*_*C*5_, is reduced to 0.26. Strains 1, 2 and 3 are kept with a growth rate of 0.27. As Figure 10 shows, the weaker strains are unable to outcompete the other three and are quickly eliminated. Changing other constants relating to the fitness of a strain achieve similar effects, the phenomenon is not unique to only altering the growth rate. Making only very small reductions to the growth rate can result in a strain surviving at a lower average population size, although this may only be due to the time needed for extinction to occur being too long to observe in these simulations.

**Fig 10.**
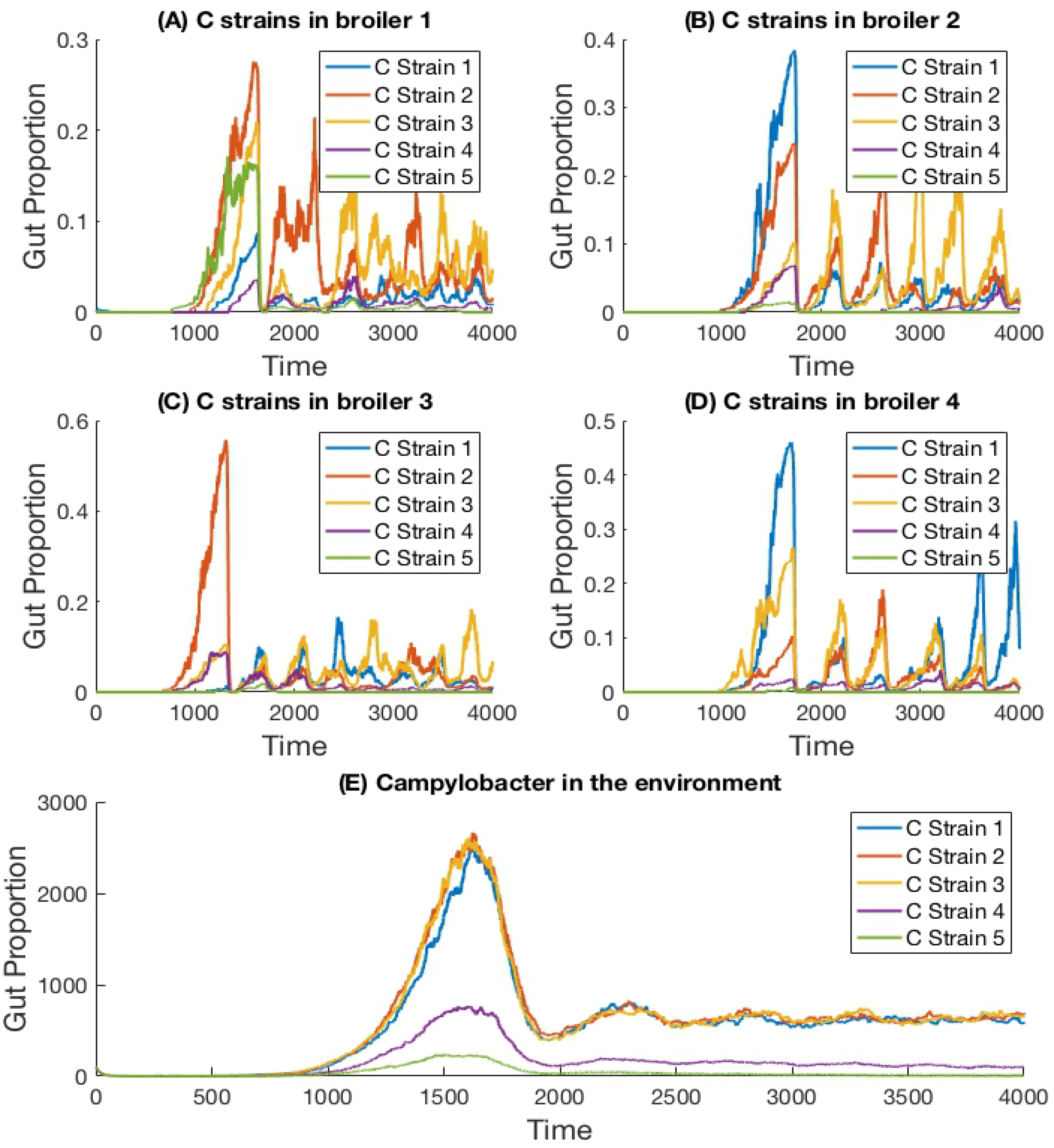
Stochastic simulations of multiple *Campylobacter* strains, differing in growth rates, within multiple broilers. The proportion of four different broilers’ microbiomes that contain five strains of *Campylobacter*. Strain 4 has had it’s growth rate reduced from 0.27 to 0.265 and strain 5 has had its growth rate reduced to 0.26. Strains 1, 2 and 3 have a growth rate of 0.27. **(E)** shows how much of the environment (total size of 200,000) contains the five strains of *Campylobacter*. These are variables *E*_*j*_ in the model.

## 3 Sensitivity Analysis

A powerful use of this model is to conduct a robust sensitivity analysis to identify the parameters of greatest impact in driving outbreaks of *Campylobacter*. We adopt a variance-based analysis of the model, and investigate the likelihood of flocks remaining free of *Campylobacter* based on a random assignment of parameter values.

We consider the case of a flock of broilers infected with a single strain of *Campylobacter*, the scenario shown in section 2.3. Model parameters are sampled randomly from a uniform range, and the model is run multiple times for these values. We then record how many of these stochastic runs resulted in the flock successfully eliminating *Campylobacter* infections, before drawing a new random sample of parameters values and repeating as necessary. Eventually we finish with a final data set which we display an example of below in Figure 11.

**Fig 11.**
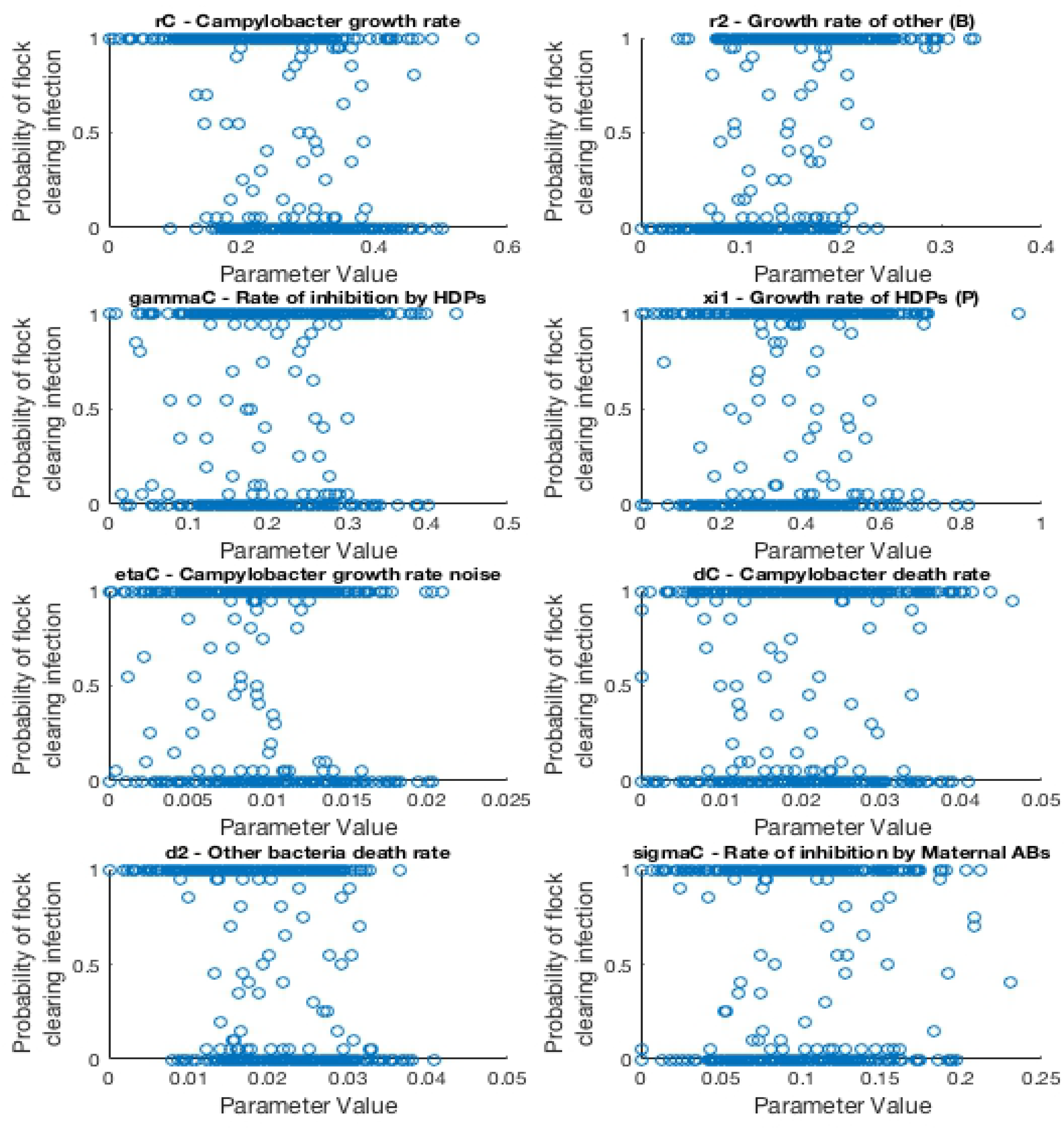
Scatter plots displaying probability of a flock clearing *Campylobacter* infection against randomly sampled parameter values. Each scatter plot depicts the results for a specific parameter value. Probability is calculated by running the model for a sampled parameter set twenty times, and recording how many of those runs resulted in the flock not becoming infected with *Campylobacter*.

As such, the most “important” parameters will be the ones which exhibit a strong trend in their scatter plot. A seemingly randomly distributed scatter plot would indicate a parameter value which has little impact on our output. To report more accurately this measure we use the first-order sensitivity index, *S*_*i*_, and the total effect index, *S*_*T*_*i*__, defined as:

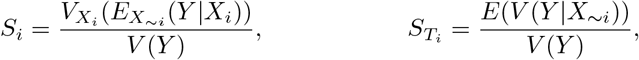

where *X*_*i*_ denotes parameter *i*, and *Y* denotes the model output. *X*_*∼i*_ denotes the vector of all factors but *X*_*i*_. *V*(*·*) denotes the variance, and *E*(*·*) the expectation. Specifically *E*(*A|B*) denotes the expectation of variable A when B is held fixed. In short *S*_*i*_ will measure the changes observed in the output when parameter *X*_*i*_ is kept fixed, while *S*_*T*_*i*__ measures the changes to the output when all other parameters are kept fixed. A full derivation and explanation can be found in Saltelli et al. (2008) [38]. In short, both are values that range from zero to one, that explain the impact of a parameter on the model output. The higher the value, the more “important” the parameter is. *S*_*T*_*i*__ is considered a stronger metric, as it also considers the higher-order impact of a parameter, whereas *S*_*i*_ only considers the immediate first-order impact. As such *S*_*i*_ would be a sufficient measure for a linear model, but for a more complex model such as the one presented in this paper, *S*_*T*_*i*__ can better reveal the impact that each parameter plays. An initial sensitivity analysis was run for twenty parameters with 1, 000 parameter set samples, drawn from a quasi-random Sobol set [38]. The results of this analysis are displayed in Table 2, and the code used to produce them is available to access at: https://osf.io/b3duc/.

**Table 2.**
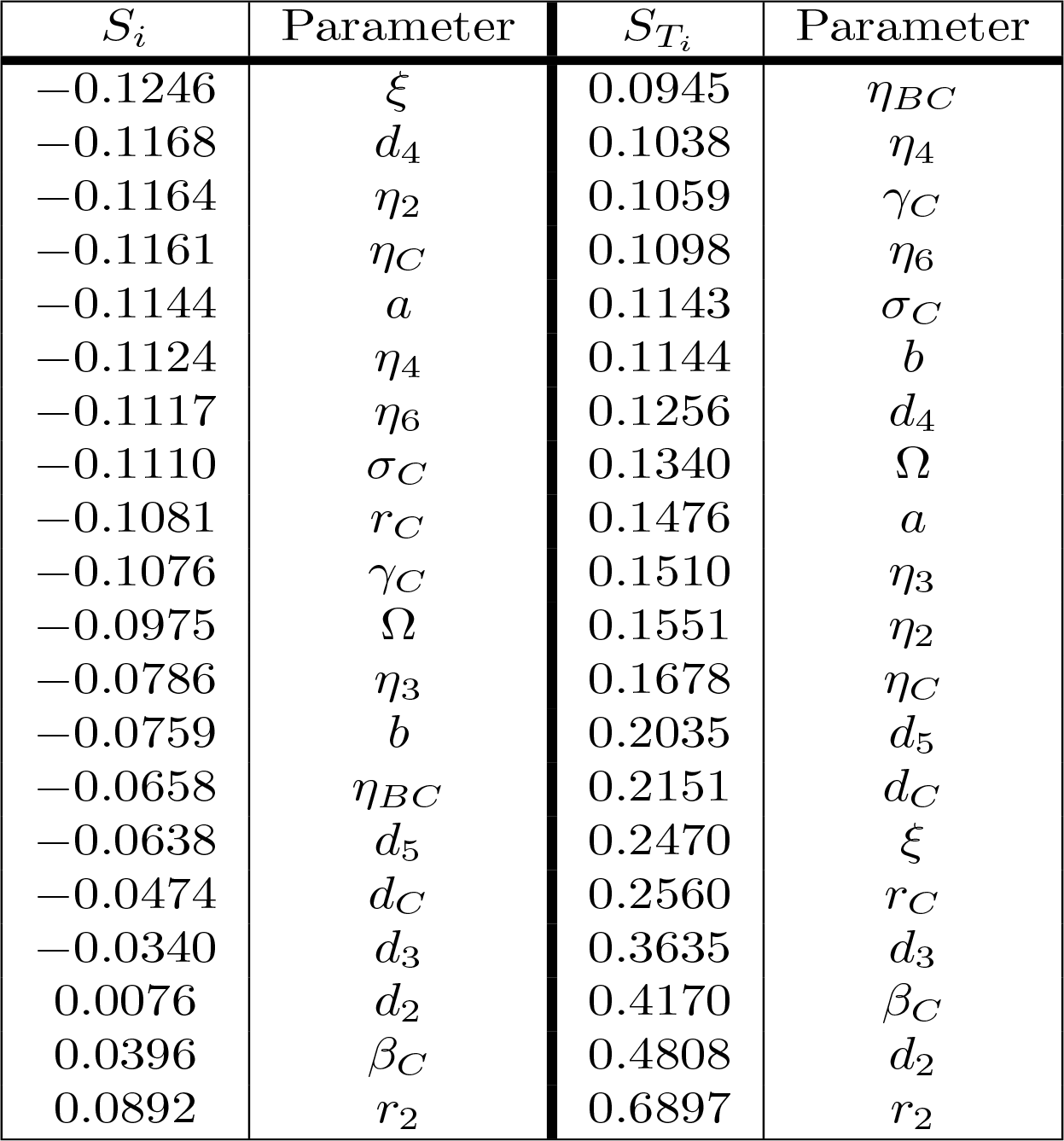
Sensitivity analysis of parameters in a stochastic model for one *Campylobacter* strain in a flock of broilers. The first-order sensitivity index and total effect index is given for a sensitivity analysis of 1, 000 runs for 20 parameters. The output function considered is the probability of *Campylobacter* going extinct within the flock based on the given parameter set.

| $S_i$ | Parameter | $S_{T_i}$ | Parameter |
| --- | --- | --- | --- |
| -0.1246 | $\xi$ | 0.0945 | $\eta_{BC}$ |
| -0.1168 | $d_4$ | 0.1038 | $\eta_4$ |
| -0.1164 | $\eta_2$ | 0.1059 | $\gamma_C$ |
| -0.1161 | $\eta_C$ | 0.1098 | $\eta_6$ |
| -0.1144 | $a$ | 0.1143 | $\sigma_C$ |
| -0.1124 | $\eta_4$ | 0.1144 | $b$ |
| -0.1117 | $\eta_6$ | 0.1256 | $d_4$ |
| -0.1110 | $\sigma_C$ | 0.1340 | $\Omega$ |
| -0.1081 | $r_C$ | 0.1476 | $a$ |
| -0.1076 | $\gamma_C$ | 0.1510 | $\eta_3$ |
| -0.0975 | $\Omega$ | 0.1551 | $\eta_2$ |
| -0.0786 | $\eta_3$ | 0.1678 | $\eta_C$ |
| -0.0759 | $b$ | 0.2035 | $d_5$ |
| -0.0658 | $\eta_{BC}$ | 0.2151 | $d_C$ |
| -0.0638 | $d_5$ | 0.2470 | $\xi$ |
| -0.0474 | $d_C$ | 0.2560 | $r_C$ |
| -0.0340 | $d_3$ | 0.3635 | $d_3$ |
| 0.0076 | $d_2$ | 0.4170 | $\beta_C$ |
| 0.0396 | $\beta_C$ | 0.4808 | $d_2$ |
| 0.0892 | $r_2$ | 0.6897 | $r_2$ |

Specifically, our objective function will run the stochastic model for a flock of chickens with the random parameter set drawn. If this model run results in no *Campylobacter* being present in the flock, it is considered to have successfully eliminated infection. The model is run twenty times with this parameter set, and the proportion of these twenty runs that results in an elimination of *Campylobacter* is the final output value, the ‘probability of flock clearing infection’.

Note that many of the *S*_*i*_ values in Table 2 are negative, despite *S*_*i*_ being limited to being between zero and one. This is due to the computational error in estimating the value, however the ordering of parameters for these particular runs will not be affected by this error. Table 2 shows that the *S*_*T*_*i*__ values associated with most parameters ranges between 0.1 and 0.2. The “most important” parameters however have a wider spread of associated *S*_*T*_*i*__ values. Stochastic simulations in particular are intensely computationally expensive, and as such, we run our sensitivity analysis a second time with a larger number of samples, using a reduced parameter set based on the initial sensitivity analysis, which we present in Table 3. We focus on the eight most important parameters from Table 2, as their sensitivity indices were highest and most varied, suggesting their impact was most distinguishable from the other parameters.

**Table 3.**
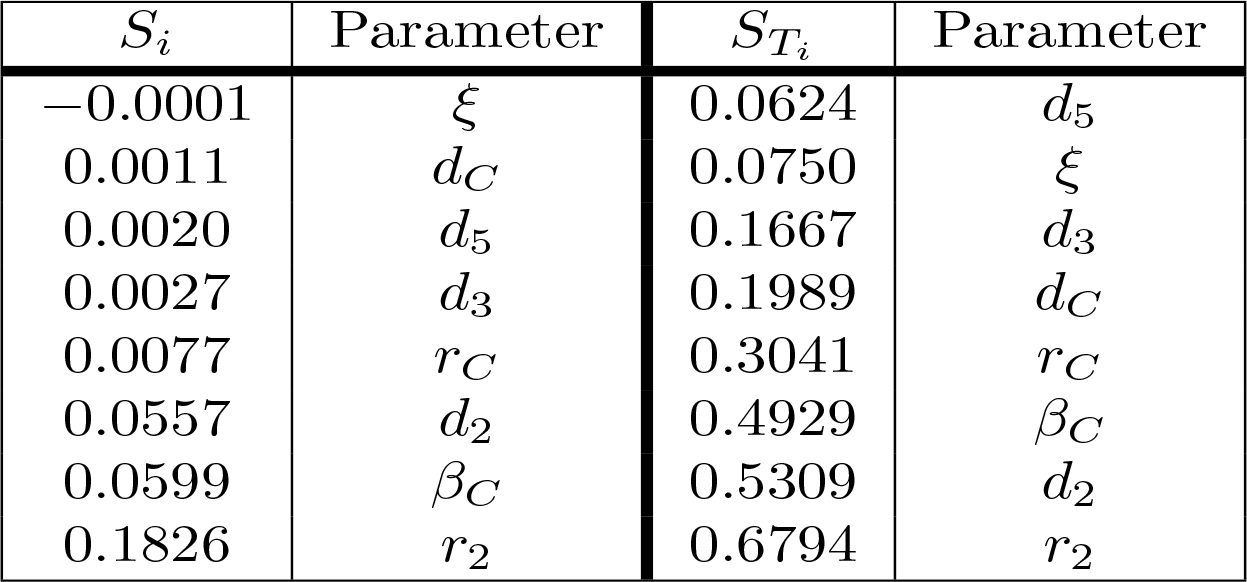
Repeated sensitivity analysis of parameters in a stochastic model for one *Campylobacter* strain in a flock of broilers. The first-order sensitivity index and total effect index is given for a sensitivity analysis of now 4; 000 runs for 8 parameters. The output function considered is the probability of *Campylobacter* going extinct within the flock based on the given parameter set.

| $S_i$ | Parameter | $S_{T_i}$ | Parameter |
| --- | --- | --- | --- |
| -0.0001 | $\xi$ | 0.0624 | $d_5$ |
| 0.0011 | $d_C$ | 0.0750 | $\xi$ |
| 0.0020 | $d_5$ | 0.1667 | $d_3$ |
| 0.0027 | $d_3$ | 0.1989 | $d_C$ |
| 0.0077 | $r_C$ | 0.3041 | $r_C$ |
| 0.0557 | $d_2$ | 0.4929 | $\beta_C$ |
| 0.0599 | $\beta_C$ | 0.5309 | $d_2$ |
| 0.1826 | $r_2$ | 0.6794 | $r_2$ |

The main result from these analyses is that the growth, death and inhibition rates of the other bacteria present in a broiler’s gut (parameters *r*_2_, *d*_2_ and *β*_*C*_) have the largest impact in eliminating *Campylobacter* from a flock. As such, we can begin to consider which preventative methods could best take advantage of this heightened sensitivity.

## 4 Discussion

Here, we have investigated the dynamics of *Campylobacter* across a range of model applications. Our framework reveals several key dynamics of microbial interaction that explain many experimentally observed phenomena. This presents promising new approaches to understanding and tackling this bacteria.

First, the most apparent prediction is that the *Campylobacter* population is successfully suppressed by the innate maternal antibodies (an experimentally observed phenomenon [39]), until these antibodies are eventually removed from the system. At this point an initial surge in the population of *Campylobacter* is observed, before it comes to rest at a lower level, reaching an equilibrium with the broiler’s immune-response. This can be seen in all of the above figures, but most clearly in Figure 1. This initial surge creates an interesting opportunity for certain strains of *Campylobacter* to emerge as an early dominating strain. Figure 5B shows that, due to the antibacterial properties of a broiler’s maternal antibodies, any strains that infect a broiler early on in its lifespan will be heavily inhibited. This creates a brief window at the point in which maternal antibodies have depleted, whereby any new strain introduced is observed to quickly colonise and dominate the gut flora, suppressing other strains (see Figure 10C). This hypothesis has been verified experimentally [39].

The proposition of damped oscillations between *Campylobacter* population size and the host’s immune-response is better reinforced by observations that host antibody populations will also oscillate in birds infected with *Campylobacter* [25]. This basic interaction has been experimentally observed by Achen et al. (1998) [29], with a high degree of variability between birds. This variability is better captured by the stochastic model, as shown in Figure 6. Indeed, many birds in Achen et al.’s study are shown to successfully clear *Campylobacter* from their system, a result rarely observed on commercial broiler farms. Likewise this result was only observed in the model case of individual, isolated broilers (see Figure 6D).

Most important is the mechanism observed in Figure 7, where the broad spectrum of oscillatory behaviour observed within a broiler is greatly reduced in a large flock of birds. Indeed the vast examples of individual dynamics observed in Figure 6, large oscilations and perhaps extinctions, are completely replaced by the same, homogenised dynamics seen within flock-reared birds in Figures 7A and 7B, as the populations of *Campylobacter* within each bird are consistently reinforced by the amount of *Campylobacter* in the environment. The wealth of experiments in monitoring flock *Campylobacter* expression for varying flock sizes means this effect can be observed taking place across multiple experiments of different flock magnitudes and densities. Morishita et al. (1997) [31] measured the amount of *Campylobacter* in a flock of thirty birds in a sizeable pen. This flock was small enough to observe oscillating behaviour in the prevalence of *Campylobacter*, and yet there do not appear to be any clear cases of birds being able to clear the bacteria from their system. Stern et al. (2001) [8] experimented with flocks of 70 birds at a density of 15.4 birds/m^2^. A small cyclic pattern is observable in their results but there are clearly far higher incidence rates. Lastly, Van Gerwe et al. (2005) [18] studied flocks of 400 birds housed at 20 birds/m^2^ (the same density considered in the above flock modelling), where now no cyclic patterns can be observed, and all birds quickly reach a constant state of *Campylobacter* expression. This effect is seen in Figure 7, and almost always observed in commercial farms [7] [40]. Our work presented here is the first, to our knowledge, to be able to propose a mechanistic explanation for this observed effect.

This dynamic, whereby broilers are consistently infected with *Campylobacter* due to highly contaminated living space, can also explain the observed phenomena whereby broiler breeder flocks (flocks kept for the breeding of meat birds) display a consistently lower *Campylobacter* prevalence rate than commercial broiler flocks [41]. Breeder birds will regularly move between periods of testing positive and negative for *Campylobacter*, inconsistently with the state of other birds in the flock, unlike the much younger birds grown for meat which remain consistently positive. Our case studies suggest that this may be due to the lower stocking density afforded to breeder birds, as it would appear the route of infection between breeder birds is weaker than that between broilers. Our sensitivity analysis however also highlighted that the gut flora can have a strong impact on the survival of *Campylobacter*. The differences in diet and management practise for breeder birds likely results in a different variety of bacterial colonies to broilers, which could also be a cause of the differences seen between breeders and broilers in *Campylobacter* expression.

Over time, our model shows strains of equal fitness will tend to settle at equal levels of prevalence on average across a flock (Figure 8E), a result that has also been shown experimentally [42] [43]. However, it is very common for an individual broiler to have only one or two dominant strains against far smaller proportions of other strains (Figures 8A - 8D and Figure 9). This effect is most prominently seen early on in the chicken’s lifespan, where usually only one strain will be present during the initial population surge of *Campylobacter*. Evidently, when one strain is already well-established within a chicken’s gut, it is difficult for a new competing strains to grow. This is due to the broiler already having a heightened level of immune response (*P*) due to the currently present strain. In the deterministic case, later strains would never be able to establish themselves as much as strains that were earlier to arrive (Figure 5B). However, in the stochastic model, there is the potential for a stochastic event to reduce the population of the currently dominating strain, and increase the population of a less-established strain.

Across the whole flock, weaker strains can be quickly out-competed by other strains. Figure 10 shows two weaker strains (strains with lower growth rates) attempting to survive within a flock, even having a slight population peak at the optimal point of strain introduction, before eventually being forced to extinction by the other three strains. Parameter variation showed that reducing a strain’s capabilities by a very small amount can allow it to persist still in the flock at a smaller average population than the others, but the majority of realisations would always end with weaker strains becoming extinct. Clearly this shows an environment where genetic dominance is very quickly selected for.

These results have considerable implications for biosecurity. While smaller flocks may have a very real opportunity to be protected from *Campylobacter* invasions, larger commercial flocks are seemingly an all-or-nothing affair. Efforts can be made to prevent initial inoculations, but once a bacterial presence is established, it may be all but impossible to remove from a flock. Considerable improvements to biosecurity have been made in recent years, but very little impact has been observed in this having reduced *Campylobacter* incidence [13]. These measures do not reduce the speed of proliferation of the bacteria, and our results suggest that better attention to bird health is likely to have a greater effect on preventing flock infection.

This model’s greatest strength is its lack of overarching assumptions. We model only the most basic bacterial interactions, all supported and verified through experimental work. Our stochastic system is capable of exhibiting a plethora of interesting dynamical interactions based on just a few known biological interactions. In moving forward with this work, the model can be used to theorise optimal methods by which to decrease the likelihood of *Campylobacter* outbreaks, and begin collaborative efforts in better explaining the evolving genetic diversity of this bacteria.

One area in which the model is admittedly lacking currently, is that it does not represent the physiological changes that occur as a bird grows. Broilers have been genetically selected over the many decades to grow excessively fast, which has been shown to have numerous concerning implications for their health [44]. This is likely to then result in differences to their auto-immune capabilities over time. More pertinently, the gut flora of a chicken is known to change and develop as the birds age [45], suggesting varying degrees of inter-bacterial uncertainty.

Our sensitivity analysis gives great insight into the optimal routes of infection prevention. Table 2 clearly shows that bolstering the growth rate and inhibition capabilities of the other bacteria populating a broiler’s gut is the best way to force extinction of *Campylobacter*, primarily through suppressing *Campylobacter* at its initial appearance in a system, before it has the opportunity to propagate. As such, the sensitivity analysis suggests further exploration and experimentation into the impact of factors which would affect the gut flora of a broiler. Probiotics are a clear way of impacting the microflora [46] and have shown some effect in studies into their impact on *Campylobacter* expression [47]. Equally, the stressors linked with stocking density have been shown to affect the gut microflora by Guardia et al. (2011) [48]. Burkholder et al. (2008) [49] have shown that feed withdrawal and heat stress can considerably alter and limit the gut microflora. These highlight that general bird health and welfare can be equally strong factors in determining the values of *r*_2_, *d*_2_ and *β*_*C*_; the parameters highlight as most “important” by the sensitivity analysis. Table 2 also however highlights the importance of parameters *ξ* and *d*_3_, the growth and death rate of host defence peptides respectively. These parameters have been shown to be strongly affected by stressors such as overcrowding [50]. As such, this result would lend further support to giving greater care to the health and welfare of broilers, as the resulting improvement to host defence peptide production would have a positive impact on helping prevent *Campylobacter* outbreaks.

These caveats notwithstanding, the model presented is capable of mechanistically explaining a wealth of experimentally observed *Campylobacter* population dynamics, further elucidating an urgent public health risk. We have used our framework to investigate multiple strain interactions, to understand better the spread of genotypes across a flock. Finally, we were able to use the model to highlight the factors most responsible for causing outbreaks of infection. Looking forward, this work can be used to understand better observed differences in outbreak dynamics between different farms and indeed countries, and further our goal of minimising public exposure to this dangerous pathogen.

